# Evolution’s boldest trick: Neurotransmission modulated whole-brain computation captures full task repertoire

**DOI:** 10.1101/2025.06.02.657368

**Authors:** Gustavo Deco, Yonatan Sanz Perl, Jakub Vohryzek, Andrea Luppi, Morten L. Kringelbach

## Abstract

The perhaps most important unsolved problem in neuroscience is how the brain survives in a complex world by performing a rich repertoire of computation on a minimal energy budget. The brain is much better at adapting to the multiplicity of stimuli and outcomes than current generations of computers, artificial neural deep learning and reservoir model architectures. Yet, at first glance the brain appears to use a fixed anatomical architecture to perform the necessary huge variety of computations. But evolution’s boldest trick is that in fact the brain’s effective connectivity is constantly being updated through neuromodulation to allow the rich repertoire of computation. Inspired by this, we created a whole-brain model using empirical neurotransmitter maps modulating the underlying local regional dynamics. This NEMO (neurotransmission modulated) whole-brain model is able to flexibly compute the full task repertoire and associated functional connectivity of the neuroimaging data from 971 healthy participants. For each individual we defined a measure of ‘brain computability’ as the fitting of the NEMO whole-brain model to all tasks performed by the individual. Importantly, brain computability correlates with both behavioural performance on individual tasks and with a general behavioural measure of intelligence. Overall, our proposed unifying NEMO framework offers a natural way to sculpt different brain dynamics in a fixed brain architecture to compute the rich repertoire of tasks required for surviving and thriving.

## Introduction

Evolution offers a deep paradox in terms of survival: How can the seemingly fixed anatomical architecture of the brain solve the multitudes of problems arising in a complex world? The brain needs to map input with output through many different transformations, that is the brain must flexibly be able to change its internal dynamics according to the mapping needed by modifying the distribution over the variables of the system during its evolution^1,2^. Each of these internal transformations is sometimes called ‘computation’ and provides the required flexibility in mapping input to output^3^. But such flexibility is not available to a fixed architecture which has fixed dynamics and therefore is only able to perform a single fixed mapping between input and output. This makes fixed architectures inappropriate for the complexity needed for survival in a complex world.

So, if the brain has a fixed architecture, how do we perform the required multitude of computations needed for survival? The answer is perhaps evolution’s boldest trick: The brain’s architecture is in fact not fixed but instead is given the necessary flexibility by way of neuromodulation^4–7^. This allows for changing the brain’s functional connectivity at the macroscale by modifying the probability of information transmission over the synaptic cleft at the microscale. At some synaptic junctions where information transfer was blocked before, the change of neuromodulation can suddenly open up for information transfer – and vice versa at other synaptic junctions. This offers exactly the mechanisms needed for flexible computation, i.e., neuromodulation can dynamically change the local internal dynamics to modify the global distribution of variables over time which allows many different mappings from input with output. In other words, the brain’s architecture is not fixed but constantly adapting by changing the internal dynamics through neuromodulation in constant feedback with neural dynamics^8^.

Evolution’s solution for adaptive computation is much better both in terms of flexibility and energy consumption than those offered by computers and artificial neural networks^9,10^. Impressively, the human brain typically uses 10-20 Watts (Joules per second)^11^ which is several orders of magnitude less than the computations performed by supercomputers which typically use around 100,000 Watts for equivalent simulations of higher functions such as chess playing^12^. This is because classic human-made computation – such as the Turing machines which underlies the Von Neumann computer architectures inside almost all current computers – is sequential in nature and designed not for optimising energy but for space and time constraints^13–15^. In contrast, the brain is performing computation in a flexible distributed way and only a small fraction of neurons and synapses are active at any instant of time, minimising the energy consumption^16,17^. On a deeper level, this fact links energy and computation to the fundamental stochastic thermodynamic non-equilibrium properties of physical systems^3,9^.

The gap in terms of computational flexibility and energy cost of Turing machines led to the rise of artificial neural networks, which offer a distributed, parallel way of computation by separating the input and output from the internal hidden neuronal layers. In particular, reservoir computing (echo state and liquid state machines) offers significant promise by using fixed architectures and a similar separation of input and output from the internal dynamical variables such as echo-state networks and liquid state machines^18–21^. In the context of neuroscience, several groups have made progress by using brain anatomy as the reservoirs^22,23^. In this sense, computation is equal to learning the linear transformations of input to the reservoir and from the reservoir to output. But this solution would require to constantly learn and re-learn the many complex input-output mappings needed. Instead, it is important to reverse-engineer the solution found in biological brains where there is no artificial separation between input and output and where computation is the dynamic transformation of the regional internal variables, modifying the global dynamics as the distribution of different brain regions over time. This trick engineered by evolution provides the brain with the necessary flexibility to have many different mappings.

Here, we were inspired by evolution’s boldest trick to implement neurotransmitter modulated whole-brain computation. Our NEMO (neurotransmission modulated) whole-brain model is able to flexibly compute the full task repertoire and associated functional connectivity of the neuroimaging data from nearly a thousand healthy participants in the Human Connectome Project (HCP)^24,25^. Flexible computation is achieved through the modulation of the regional local dynamics in the actual fixed neuroanatomical architecture of the human brain which gives rise to multiple dynamical mappings. This ‘brain computability’ is defined for each of the individuals by fitting of the NEMO whole-brain model to all the tasks performed by the individual. Crucially, brain computability correlates with not only the behavioural performance on the different tasks but also a general behavioural measure of intelligence across individuals. Brain computability is achieved through a dynamic reconfiguration of the hierarchy of orchestration which is shown to highly correlate with the levels of thermodynamic non-equilibrium. As such, this provides a key link between computation, brain dynamics and thermodynamic non-equilibrium. The link with thermodynamic non-equilibrium is essential to the present endeavour because there is a deep connection between thermodynamic work (which implies energy consumption), information and computation, as introduced in the famous example of Maxwell’s Demon^3,9^. Overall, the NEMO framework demonstrates the powerful utility of evolution’s boldest trick in sculpting different brain dynamics in a seemingly fixed brain architecture to compute the rich repertoire of tasks required for survival.

## Results

We investigated the explanatory power of neurotransmission modulated (NEMO) whole-brain computation in a large-scale dataset of 971 participants from the Human Connectome Project (HCP) during resting state and while performing seven different tasks [EMOTION, RELATIONAL, GAMBLING, SOCIAL, LANGUAGE, MOTOR, WM (working memory)], designed to cover most of the cognitive and emotional domain. The key idea is to show how a fixed brain architecture becomes flexible through neuromodulation in such a way that the dynamics change to be able to perform the multiple different computations required for each of the tasks.

The overall scheme is summarised in **Figure 1**. Starting with the history of reservoir computing, **Figure 1A** shows how artificial neural networks perform computation by mapping input to output through the dynamics of a fixed, non-linear reservoir, whether consisting of a recurrent artificial neural network (top) or an empirical brain connectivity network (bottom). The core idea of reservoir computing is that the reservoir maps input signals into a higher dimensional computational space which through a readout mechanism is in turn mapped to output. Interestingly, the required maps between input and reservoir, as well as between reservoir and output are linear transformations. While this approach is universal and powerful, different computations require a remapping of the fixed reservoir with different input and output signals which is very unlike the brain which has no time for such relearning but instead operates in a time-critical, on-the-fly manner. Therefore, reservoir computing with a fixed architecture is an imperfect simulation of how the biological brain achieves its remarkable flexibility in the face of real life problems.

**Figure 1.**
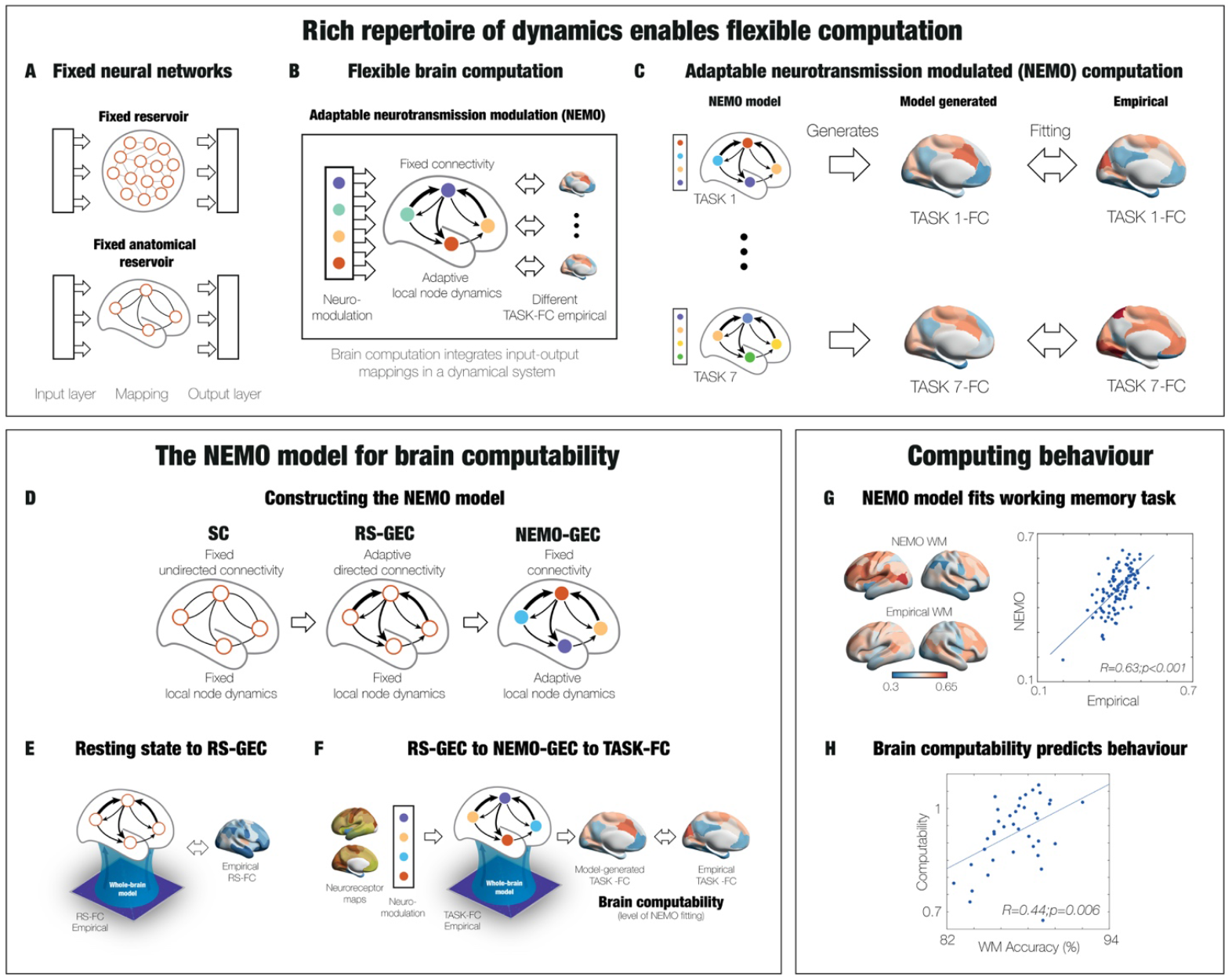
Neurotransmission modulated (NEMO) whole-brain computation. **A)** Current generations of artificial neural networks typically perform computation by mapping input to output through hidden layers or dynamical reservoirs. Classic reservoir computing (top) creates a general transformation between input and output using a fixed but rich reservoir. One way of making the reservoir more similar to the workings of the brain is to use the actual empirical anatomical (undirected) connectivity as the reservoir (bottom). However, this fixed architecture is unable to accommodate the required multitude of computations needed for survival. **B)** Instead, the brain achieves flexible computation through the bold trick of evolution, namely through dynamic neuromodulation of the local internal dynamics that modify the global distribution of variables over time allowing for many different mappings from input with output. The figure shows the implementation of the NEMO whole-brain model, which in a similar way uses neurotransmission maps to update the regional weights (the coloured circles). **C)** The local dynamics in the fixed, directed connectivity generate global dynamics that can be flexibly adapted to fit any task (here the seven HCP tasks). **D)** The panel shows how the NEMO model is created from the initial structural connectivity (SC, left) which has fixed undirected connectivity to RS-GEC (middle) and to NEMO-GEC (right). **E)** The SC is used to fit the resting state whole-brain model of an individual, which generates the resting-state generative effective connectivity (RS-GEC, middle). This resting state whole-brain model contains directed connectivity with fixed, homogeneous local node-dynamics. **F)** Moving beyond this to do actual brain computation, NEMO-GEC is generated by adapting the neuromodulation to fit the functional connectivity of each of the seven HCP tasks by changing only the local dynamics on the RS-GEC model (but preserving the connectivity). We define brain computability for each individual as the fitting of the NEMO whole-brain model to all the tasks performed. **G)** Using NEMO on tasks provides excellent individual fits, for example in the working memory task. **H)** Equally, the brain computability significantly captures behaviour such as task performance.

In contrast, as shown in **Figure 1B**, here we propose to reuse evolution’s bold trick of achieving flexible computation through dynamic neuromodulation of the local internal dynamics. In other words, the underlying connectivity stays fixed, while the regional dynamics are changed through neuromodulation in such a way that the global dynamics is hierarchically reconfigured, flexibly allowing for many different computations. As such brain computation can be defined as the flexible change of the system’s internal global dynamics according to the mapping needed, by modifying the distribution over the regional variables of the system over time. The figure shows the implementation of these principles in the NEMO whole-brain model using neurotransmission maps to update the regional weights (indicated by the differently coloured circles) to change the local dynamics.

**Figure 1C** shows the process through which the local dynamics in the fixed, directed connectivity in the NEMO whole-brain model reconfigure the hierarchy of the global dynamics so that this can sustain the different computations for each of the seven HCP tasks.

Here, we provide a summary of how the NEMO whole-brain model is constructed but the details are found in the Methods. First, this is based on the principles of whole-brain modelling, which combines the anatomical connectivity with local dynamics to fit the dynamics of empirical neuroimaging data^26–28^. The local dynamics can be simulated using many different models such as spiking, dynamical mean field and Hopf local regional models for fitting to the empirical neuroimaging data. Here, we chose the Hopf model to fit the neuroimaging data, since this model has been shown to provide the best fit to empirical data^29–32^.

As shown in **Figure 1D**, there are three steps needed to construct the NEMO whole-brain architecture: SC (structural connectivity), RS-GEC (resting state generative effective connectivity) and NEMO-GEC.

The first step (shown in **Figure 1E**) is to use the structural, fixed undirected connectivity (SC, left) reconstructed from in vivo diffusion tractography to start fitting the empirical resting state (RS) fMRI data of an individual. Then, as the second step, the model fitting process iteratively adjust a generative effective connectivity (GEC), which is given as asymmetric weights of the existing anatomical connections^30^ and extends the concept of effective connectivity^33^. However, GEC provides a generative measure of the whole-brain network because it uses the whole-brain model to adapt the strength of existing anatomical connectivity, i.e., the effective conductivity values of each fibre (see Methods for details). This generates a RS-GEC whole-brain model of the resting state with RS-GEC connectivity (middle), i.e., directed connectivity with fixed, homogeneous local node-dynamics.

The third step, shown in **Figure 1F**, is to generate a NEMO whole-brain model for actual brain computation needed for each task. In an iterative process (described in detail in the Methods), NEMO-GEC is generated by adapting the neuromodulation to fit the fMRI functional connectivity of each of the seven HCP tasks. This is done by preserving the connectivity but modulating only the local dynamics of the RS-GEC model. As such the *brain computability* is defined for each individual as the fitting of the NEMO whole-brain model to the functional connectivity of all seven tasks.

The figure also shows examples of the results of using NEMO on tasks which provide excellent individual fits to for example the working memory task (**Figure 1G**) and how the brain computability significantly captures behaviour such as task performance (**Figure 1H**).

### Principles of brain computation through neurotransmitter modulation

The neuromodulation of the NEMO whole-brain model relies on using empirical neurotransmitter maps. **Figure 2A** shows the rendering in the Schaefer100 parcellation of 19 different neurotransmitter receptors and transporters across nine different neurotransmitter systems (dopamine, serotonin, norepinephrine, glutamate, acetylcholine, GABA, histamine, cannabinoid and opioid). These maps used PET tracer images from more than 1,200 healthy participants which were collated and averaged to produce mean receptor distribution maps^34^.

**Figure 2.**
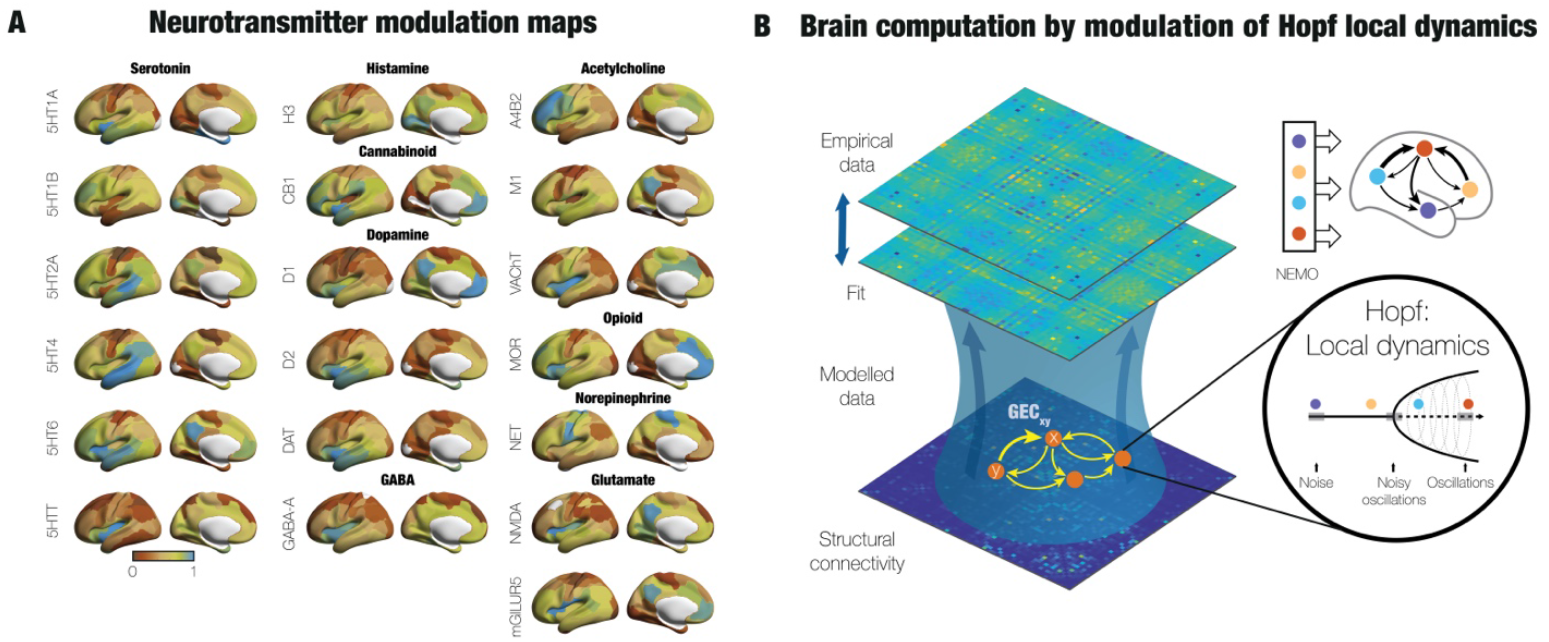
Principles of brain computation through neurotransmitter modulation. **A)** The panel shows 19 different neurotransmitter receptors and transporters across nine different neurotransmitter systems. These maps used PET tracer images from more than 1,200 healthy participants which were collated and averaged to produce mean receptor distribution maps^34^. **B)** Whole-brain models link anatomy structure with function by fitting the empirical neuroimaging resting-state data in a given parcellation. In each region the local dynamics have traditionally been governed by a homogeneous Hopf bifurcation model (Stuart-Landau oscillators) coupled through the brain’s structural connectivity. The conductivity of the existing anatomical connections is optimised to fit the whole-brain model to a global measure of the empirical neuroimaging data, typically the functional connectivity matrix. This optimisation gives rise to the model’s resting-state generative effective connectivity (RS-GEC), which is a directed matrix of connectivity masked by the existing anatomy. In order for this RS-GEC model to be able to model different tasks, we introduced local heterogeneity by changing the noise level of the Hopf model according to neuromodulation in each region. Given that the local dynamics of the Hopf model is working at the edge of the bifurcation, the modulation by noise can change the local dynamics to become more noisy or more oscillatory. In turn, this changes global dynamics without changing the underlying RS-GEC fixed architecture. This is exactly what is needed to perform multiple brain computation in a flexible way.

These neurotransmitter maps are then used to change the local Hopf model coupled through the resting-state generative effective connectivity (RS-GEC) obtained for each individual in the way described above (**Figure 2B**). We change the local heterogeneity by changing the noise level of each local Hopf model in the parcellation according to the weighted neuromodulation. The modulation by noise can change the local dynamics to become more noisy or more oscillatory, since the local dynamics of the Hopf model is working at the edge of the critical bifurcation between noisy and oscillatory regimes^26,35,36^. These local changes reconfigure global dynamics to perform multiple brain computation in a flexible way for each task, but importantly without changing the underlying fixed architecture. Please note that the local regional changes all use the same optimised weighted linear combination of the heterogenous 19 neurotransmitter maps.

In order for the NEMO model to be able to model the seven different tasks, we used the high dimensional swarm optimisation algorithm (see *Methods*) to find the optimal weighting of the 19 neurotransmitters across the full parcellation to fit the empirical functional connectivity of each task. The swarm algorithm traverses the parameter search domain similarly to how a swarm of birds would explore a space and has been shown to be a robust tool for global optimisation of higher dimensional space. Here, the swarm algorithm used the 19 neurotransmitter weighting parameters as well as two free parameters adapting the extrema (see *Methods*).

### Brain computation by neurotransmission modulation (NEMO) whole-brain model

The swarm optimisation produced the optimal linear weightings of the 19 neurotransmitter maps fitting the empirical functional connectivity of each of the seven HCP tasks. **Figure 3A** shows the results as a matrix with each of the seven HCP tasks versus the individual weighted contribution of the neurotransmitters from −1 to 1. For each task, these weights then multiply each of the individual normalised 19 neurotransmitter maps (see *Methods*) to produce a final weighted map of neuromodulation of the Hopf model in each region of the Schaefer100 parcellation. This weighting of the 19 neurotransmitter spatial maps give rise to the different noise levels for the regions in each of the seven tasks (see **Equations 9** and **10** in the *Methods*).

**Figure 3.**
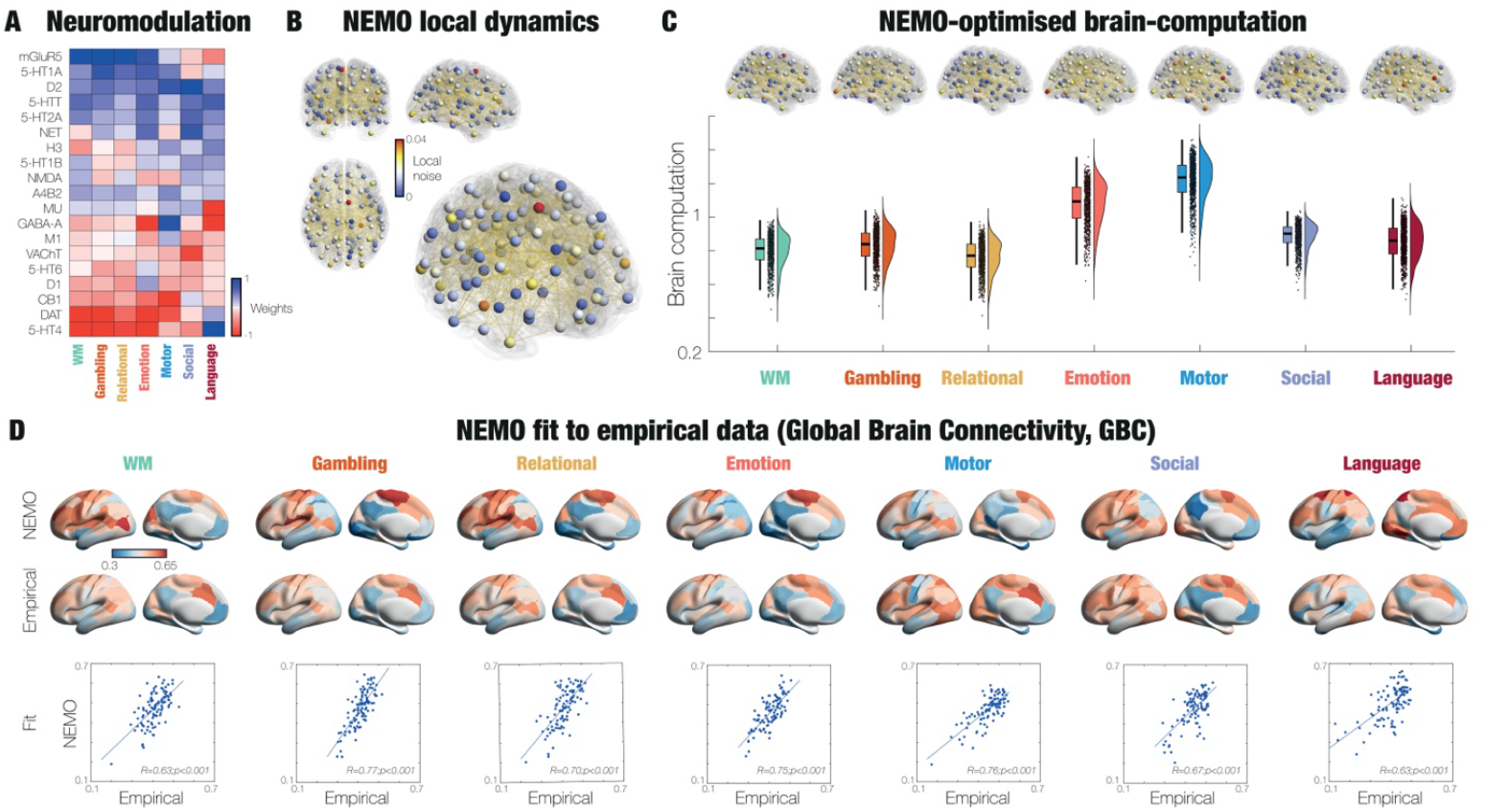
Neurotransmission modulation (NEMO) needed for brain computation. **A)** The matrix shows the weighted contribution of each of the 19 neurotransmitter systems for the seven HCP tasks after optimisation with NEMO whole-brain model. Take for example the Gambling task, where the first four neurotransmitters listed (mGLUR5, 5-HT1A, D2 and 5-HTT) have the highest positive weights, while the last two neurotransmitter (DAT and 5-HT4) have the highest negative weights. This means that the glutamate, dopamine and serotonergic neurotransmission contribute most to the brain computation involved in gambling, which is consistent with the literature^37,38^. **B)** The rendering is showing the 100 local regions in the Schaefer100 parcellation with each region coloured according to the optimal noise level coming from the weighted neuromodulation for the working memory task. The clockwise renderings (from top left) are showing the brain from the front, side, 3D and top. **C)** The boxplot shows the high levels of brain computability for each task, which shows that NEMO is able to capture the shift from resting state to task for all tasks. Above each of the boxplots, there are renderings of the optimal noise level for each task (similar to panel B). Interestingly, the best level of brain computability is found for Motor, which is of course in many ways as different as possible from resting state. **D)** Similarly, the high level of fitting to tasks is demonstrated by the very similar renderings (lateral and medial views) global brain connectivity (GBC) for the Schaefer100 parcellation in the model and in empirical data (top). The scatterplots (bottom) for each of the tasks show significant correlations between the average regional GBC across the 971 HCP participants for all tasks.

**Figure 3B** shows four renderings of the noise map for the WM task with the RS-GEC connectivity (front, side, 3D and top views, clockwise from top left). These renderings show the ‘engine’ driving the brain computation and is an actual version of the cartoon shown in **Figures 1** and **2. Figure 3C** shows the side rendering of the ‘engine’ for every task. Please note how the different colours in each region across tasks signifies the reconfiguration of NEMO to carry out the necessary computation. Below these renderings, for each task we show a boxplot of the level of NEMO-optimised brain computation (based on data from all 971 participants). The resulting high levels of brain computation for each task show the evidence for the ability of NEMO to perform the necessary task-specific computations. As an example, the optimisation of neurotransmitters for the GAMBLING task show that the glutamate, dopamine and serotonergic neurotransmission contribute most to the brain computation involved in gambling, which is consistent with the large literature^37,38^.

The fitting of the NEMO model to empirical functional connectivity is shown in **Figure 3D**, where there are very similar renderings (lateral and medial views) of global brain connectivity (GBC) for the model and in empirical data (top). This is further shown in the scatterplots below for each of the tasks with significant correlations between the average regional GBC across the 971 HCP participants for all tasks (*R*=0.70 +/- 0.06, *p*<0.001).

### Reconfiguration of hierarchy for task computation and associated behaviour

The anatomy of the human brain is hierarchically organised which facilitates the computation of sensory input followed by higher-level computation in nested recursive circuits at various spatiotemporal levels^39–41^. Yet, neurotransmission allows for the flexible reorganisation of this hierarchy^6,7,42^. Ultimately, this flexible hierarchical organisation of the brain is necessary for the execution of time-critical computations^43–45^.

Here, we wanted to demonstrate the empirical hierarchical changes in brain organisation needed for fitting the empirical task data and explaining the behaviour. For this purpose, we generated a TASK-GEC for each task. Briefly describing the method, for each task we fitted a whole-brain model to the generated timeseries from the NEMO-GEC model. From the TASK-GEC model, we used the measure of trophic coherence to generate the hierarchical reorganisation^46^. As shown in the Methods, this method is based on previous work in ecology^47^, which has been extended to general directed networks^46^. Briefly, for a directed network, this provides both the hierarchical, node-level information, *trophic level*, a measure of where a node sits in the hierarchy of a directed network; and the global information, *directedness* (or trophic coherence). Intuitively this can seen in ecology, where low trophic levels are assigned to plants, while high trophic level nodes are assigned to carnivores, with energy flowing up the food web from low to high trophic levels. The hierarchical reorganisation in the working memory (WM) task can clearly be seen in **Figure 4A**, which shows the renderings and boxplot of hierarchy between the RS-GEC and TASK-GEC for WM. The hierarchy is significantly higher for the WM task than for resting state (p<0.001, Wilcoxon), reflecting the more directed information flow in task computation. The brain computation for each individual performing the WM task is shown in the four scatterplots against the task accuracy (top) and reaction times (bottom) for each individual (left) and for groups of 20 similar participants (right). Here we used the group-average to increase the robustness. The groups were constructed by grouping together individuals with similar task accuracy. As can be seen, we found significant correlations (*R*=0.44, *p*=0.006 for task accuracy; *R*=-0.49, *p*=0.005 for tasks reaction times).

**Figure 4.**
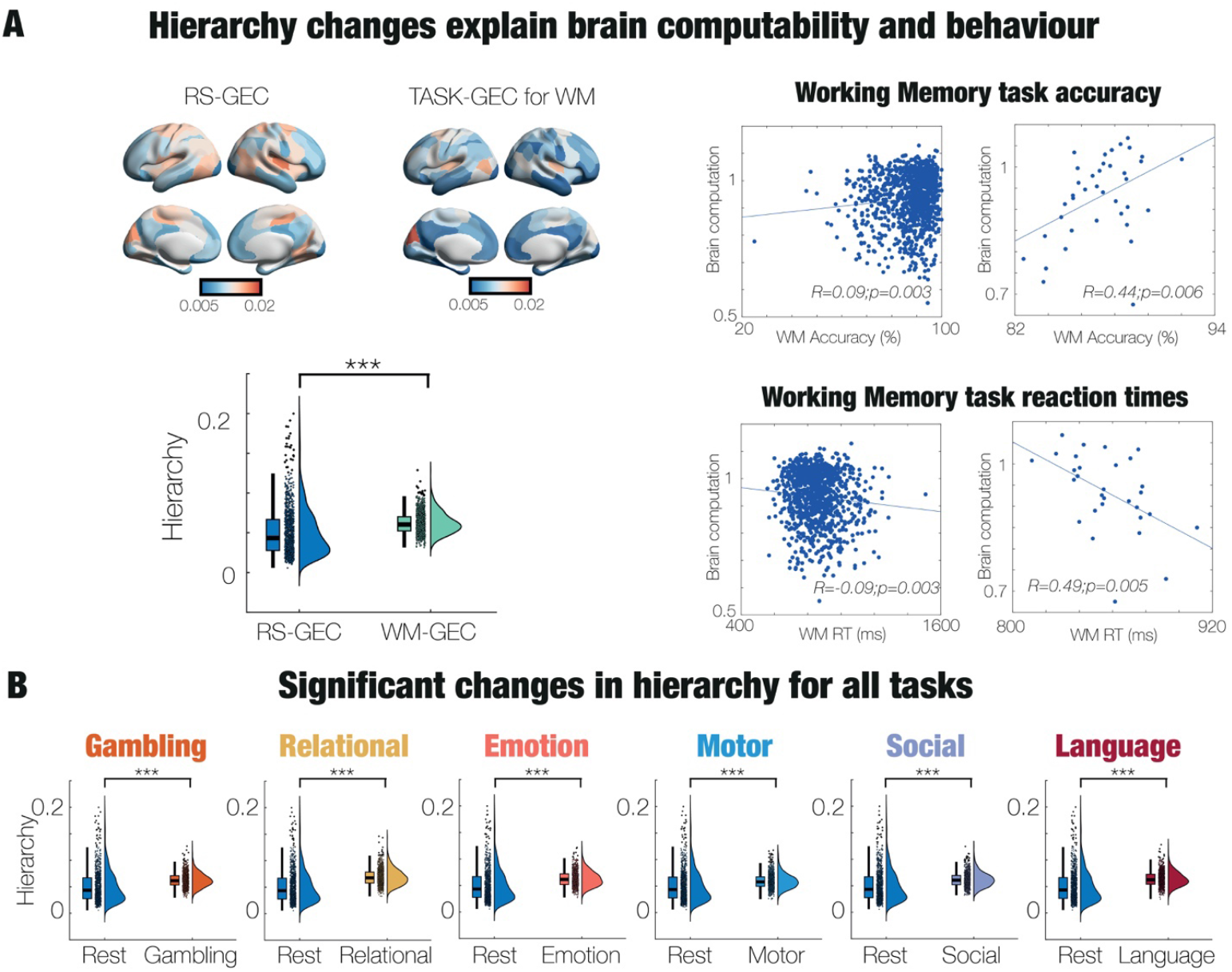
Changes in hierarchy explain task computation and associated behaviour. In order to quantify the hierarchical changes in brain organisation needed for fitting the empirical task data and explaining the behaviour, we generated a TASK-GEC for each task, starting from the NEMO-GEC. For each task, we fitted a whole-brain model to the generated timeseries from the NEMO-GEC model. From this model, trophic coherence will generate the hierarchical reorganisation for this TASK-GEC. **A)** For the working memory (WM) task the hierarchical reorganisation can clearly be seen in the different renderings and boxplot of hierarchy between the RS-GEC and TASK-GEC for WM. As can be seen, the hierarchy is significantly higher for the WM task than for resting state (p<0.001, Wilcoxon), meaning that there is more directed information flow in task computation. This difference, i.e., the brain computability for each individual performing the WM task is shown in the scatterplots against the task accuracy (top) and reaction times (bottom). The scatterplots on the left show the brain computability for each individual and for groups of 20 similar participants on the right. In both cases, there are significant correlations (see panels). B) There are significant differences in hierarchical organisation for the TASK-GEC for each task compared to the RS-GEC. The increases in hierarchy for all tasks indicate that there is more directed information flow needed for the necessary task computation.

Similarly, all the tasks have significant differences in hierarchical organisation compared to the RS-GEC. Again, the hierarchical increase for all tasks is indicative of more directed information flow needed for the necessary task computation.

### Linking brain computability to intelligence, thermodynamic non-equilibrium and brain hierarchy

As indicated by its name, brain computability measures the ability to perform computation and as such this concept must be linked to many important measures of brain performance. Here, shown in **Figure 5A** we first compared the maximal brain computability to the working point of the NEMO model, where a value of the global coupling of 1 is fitting the empirical data (see Methods). As can be seen from the figure, on average the brain dynamics in people are close to optimal brain computability

**Figure 5.**
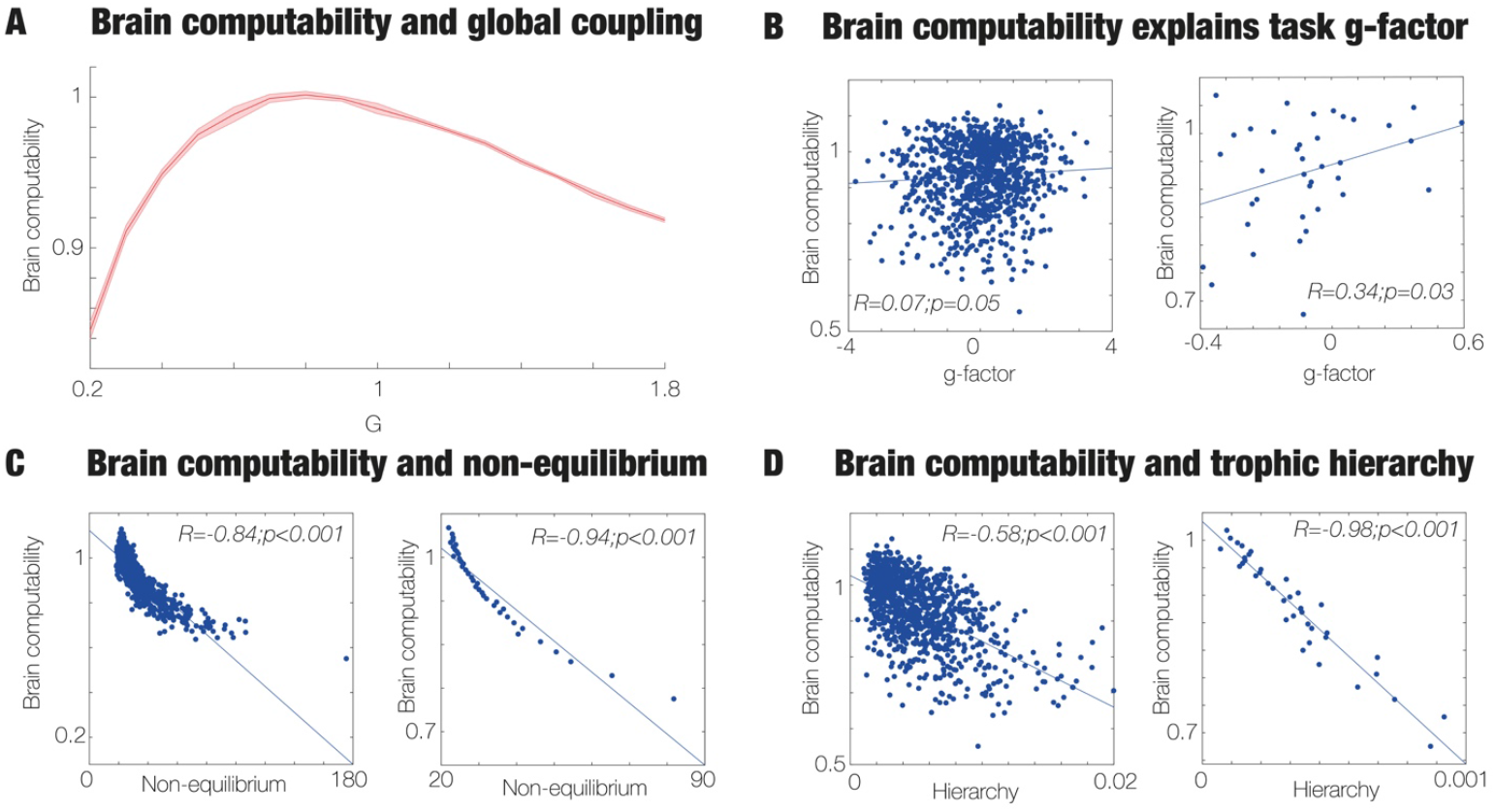
Linking brain computability, intelligence, thermodynamic non-equilibrium and brain hierarchy. **A)** The maximal brain computability is close to the working point of the NEMO model fitted to the resting state and seven tasks of the large empirical HCP neuroimaging cohort. This suggests that on average the brain dynamics in people are close to optimal brain computability. **B)** In fact, there is a significant correlation between brain computability and g-factor, a derivation of a general factor of intelligence obtained by combining measures from neuropsychological battery. The left scatterplot shows all the 971 individuals, while each dot corresponds to groups of 20 in the left scatterplot. **C)** There is also a significant negative correlation between brain computability and thermodynamic non-equilibrium, as shown in the two scatterplots (all participants on the left, groups on the right). High brain computability is linked to lower levels of non-equilibrium, while lower levels of brain computability is linked to more non-equilibrium. **D)** Similar negative correlations are seen between brain computability and brain hierarchy (measured with trophic coherence). This strongly links brain computability with intelligence, thermodynamic non-equilibrium and brain hierarchy.

This is important since **Figure 5B** shows a significant correlation between brain computability and *g*-factor, which is a general factor of intelligence obtained by combining measures from neuropsychological battery. The left scatterplot shows data from all the 971 individuals (*R*=0.07, *p*<0.05), whereas in the right scatterplot each dot corresponds to groups of 20 participants (*R*=0.34, *p*<0.03). The groups were constructed by grouping together the participants with similar brain computability.

Equally, **Figure 5C** shows that is also a significant correlation between brain computability and thermodynamic non-equilibrium, albeit a negative rather than positive correlation (*R*=-0.84, *p*<0.001 and *R*=-0.94, *p*<0.001 for the group). The two scatterplots again all participants on the left with groups of 20 on the right.

Similar, **Figure 5D** shows negative correlations between brain computability and brain hierarchy (as measured with trophic coherence, *R*=-0.58, *p*<0.001 and *R*=-0.98, *p*<0.001 for the group). This strongly links brain computability with intelligence, thermodynamic non-equilibrium and brain hierarchy.

Finally, there are also significant correlations (not shown in the figure) between non-equilibrium and hierarchy at both the individual and group level (*R*=0.55, *p*<0.001 and *R*=0.93, *p*<0.001, respectively). There were also significant correlations at the group level between g-factor and hierarchy (*R*=-0.39, *p*=0.02) as well as g-factor and non-equilibrium (*R*=-0.32, *p*=0.04).

## Discussion

Here we have presented the first neurotransmission modulated (NEMO) whole-brain model that can dynamically change the hierarchical processing needed for efficient task computation. This is based on evolution’s boldest trick of taking a seemingly fixed architecture and making it dynamic through neurotransmission. This opens up avenues for far more efficient computation in terms of energy and time compared to current approaches to computation in computers and artificial neural networks.

We demonstrated that the NEMO whole-brain model can perform multiple different task computation through using the optimal weighted sum of using 19 neurotransmitter maps to modulate the regional dynamics of the many Hopf local models driving the global activity (**Figures 2** and **3**). Importantly, the brain computability in the NEMO framework is achieved using a fixed architecture without changing the whole-brain coupling strengths. Specifically, we show that the NEMO whole-brain achieves this by changing the initial hierarchy of information flow of the resting state in each participant to a changed hierarchy for each of the seven HCP tasks for each of the 971 healthy individuals (**Figure 4**). This is reflected in the significant change in the NEMO fit from resting state to the functional connectivity of each of the tasks (**Figure 3C**).

Furthermore, attesting to the scientific strength of the NEMO framework, we can define a measure of ‘brain computability’ for each individual as the level of fitting of the NEMO whole-brain model to resting state and seven tasks. This brain computability significantly correlates with behavioural measures such as task accuracy and reaction times for the working memory task across all 971 individuals (**Figure 4A**).

The concept of ‘brain computability’ turns out to be an excellent description of the state dynamics and abilities of the brain for an individual both in resting state and when performing all seven tasks. Interestingly, on average the optimal working point of the whole-brain model to resting state is close but not identical to optimal brain computability across people (**Figure 5A**). This is an important finding since the results suggest that different people may be more or less optimal in solving tasks – which is what behavioural data indicate. In fact, this was confirmed by our finding of a significant correlation between brain computability and the *g*-factor, which is derived from a neuropsychological battery and thought to reflect general intelligence (**Figure 5B**)^48^. While often debated, the *g*-factor is thought to target important skills like verbal fluency and fluid intelligence, which is commonly taken to describe abilities related to problem solving independently of existing knowledge^49^.

We further investigated this important correlation and found that brain computability is also significantly correlated with both thermodynamical non-equilibrium and hierarchy (**Figure 5CD**). This confirms the deep link found in stochastic thermodynamics, where computation, information and energy are closely related^3,9^. This deep link to non-equilibrium thermodynamics is important since this is likely to be an especially promising way of understanding brain computation. We have previously shown that thermodynamics can be used to quantify brain hierarchy through estimating the asymmetry of information flow which provides hierarchical organisation that facilitates fast information transfer and processing at the lowest possible metabolic cost^30,32,50^. Precisely the asymmetry of information flow, which is in thermodynamics is called the ‘breaking of the detailed balance’, creates a non-equilibrium system.

In terms of brain function and in particular the ability to allow for the efficient transfer of energy/information over space and time, it has been shown that turbulence is a fundamental and highly useful non-equilibrium thermodynamic principle providing optimal mixing properties^51^. The brain was recently shown to be turbulent, where the scale-free nature of turbulence provides exactly the kind of dynamical regime, where hierarchical information cascades can provide optimality of brain function^52–54^. As such, turbulence is also a highly sensitive biomarker of different brain states such as psychedelics^55^, depression^56^ and sleep, coma and meditation^57^. Remarkably, the level of turbulence pre-treatment was also predictive of the treatment outcome of the pharmacological intervention for depression^56^.

The non-equilibrium turbulent behaviour of the brain strongly influences the evolution of the probability distribution of brain states, and thus create a strong link to the brain computability provided here. Interestingly, similar to maximum brain computability, turbulence is also slightly shifted to the left compared to the average optimal working point of the whole-brain model fit to empirical data. This suggests a link between turbulence and brain computability. An interesting hypothesis following from this could be that turbulence plays a key role in maximising computation while minimising energy consumption, reflected in the power laws found in brain turbulence^58–60^. This should be further investigated in future.

Overall, the NEMO framework is a first step to provide the necessary flexibility for the overwhelming computations needed for the brain to survive and thrive in the world. On a deeper level, the findings link computation and information to the fundamental stochastic thermodynamic non-equilibrium properties of physical systems. This shows how information and computation are entangled in the brain, which may be why less energy is used. Our findings also suggest how hierarchy and the sparsity of the GEC helps to minimise the energy consumption by limiting the information and computation. Going forward, the NEMO framework can be used to quantify the energy used for brain computability by relating computability and thermodynamic non-equilibrium, and potentially provides a road map to create true brain-like, energy-efficient artificial intelligence.

## Methods

### Human Connectome Project: Acquisition and pre-processing Ethics

The Human Connectome project is based on ethics obtained by the Washington University–University of Minnesota (WU-Minn HCP) Consortium who obtained full informed consent from all participants. Research procedures and ethical guidelines were followed in accordance with Washington University institutional review board approval (Mapping the Human Connectome: Structure, Function, and Heritability; IRB # 201204036).

### Participants

We chose a sample of 971 participants, all of whom have resting state data and performed all seven tasks. These participants were selected from the March 2017 public data release from the Human Connectome Project (HCP).

### The HCP task battery of seven tasks

All participants performed the standard HCP task battery described in detail on the HCP website ^61^. This consists of seven tasks: Working memory (WM), motor, gambling, language, social, emotional, relational. According to HCP, the tasks were designed to cover a broad range of human abilities in several major domains sampling the diversity of the brain: 1) Visual, motion, somatosensory, and motor systems, 2) category-specific representations; 3) working memory, decision-making and cognitive control systems; 4) language processing; 5) relational processing; 6) emotion processing; and 7) social cognition. All HCP participants performed all tasks in two separate sessions (first session: gambling, WM and motor; second session: language, emotion processing, relational processing and social cognition).

### 3T structural data

The HCP structural data were acquired using a customized 3 Tesla Siemens Connectom Skyra scanner with a standard Siemens 32-channel RF-receive head coil. For each participant, at least one 3D T1w MPRAGE image and one 3D T2w SPACE image were collected at 0.7 mm isotropic resolution.

### 3T diffusion MRI

In order to reconstruct a high-quality average structural connectivity (SC) matrix for constructing the whole-brain model (using the Schaefer100 parcellation), we obtained multi-shell diffusion-weighted imaging data from 32 participants from the HCP database (scanned for approximately 89 minutes). The acquisition parameters are described in detail on the HCP website^62^. The connectivity was estimated using the method described by Horn and colleagues^63^. In summary, the data was processed using a generalized q-sampling imaging algorithm implemented in DSI studio (http://dsi-studio.labsolver.org). A white-matter mask was produced from segmentation of the T2-weighted anatomical images, co-registered the images to the b0 image of the diffusion data using SPM12. For each of the 32 HCP participants, 200,000 fibres were sampled within the white-matter mask. Fibres were transformed into MNI space using Lead-DBS^64^. The methods used the algorithms for false-positive fibres shown to be optimal in recent open challenges^65,66^. Accordingly, the risk of false positive tractography was reduced by using the tracking method achieving the highest (92%) valid connection score among 96 methods submitted from 20 different research groups in a recent open competition^65^.

### Neuroimaging acquisition for fMRI HCP

All the 971 HCP participants were scanned on a 3-T connectome-Skyra scanner (Siemens). For the resting state data, we used one fMRI acquisition of approximately 15 minutes acquired on the first day, where participants had eyes open with relaxed fixation on a projected bright cross-hair on a dark background. We also used fMRI data from all the seven tasks. Full details of participants, the acquisition protocol and pre-processing of the data for both resting state and the seven tasks are provided by the HCP website (http://www.humanconnectome.org/) but here we give a brief summary.

We used the standard minimal pre-processing of the HCP resting state and task datasets (as described in full details on the HCP website). This uses the standard HCP pipeline with standardized methods including FSL (FMRIB Software Library), FreeSurfer, and the Connectome Workbench software^67,68^. This pre-processing includes correction for spatial and gradient distortions and head motion, intensity normalization and bias field removal, registration to the T1 weighted structural image, transformation to the 2mm Montreal Neurological Institute (MNI) space, and using the FIX artefact removal procedure^68,69^. Head motion parameters were regressed out and structured artefacts were removed by Independent Component Analysis followed by FMRIB’s ICA-based X-noiseifier^70,71^.

The final pre-processed timeseries are in HCP CIFTI greyordinates standard space and available in the same standard surface-based CIFTI file for each participants for resting state and each of the seven tasks.

From these pre-processed timeseries, we extracted the average timeseries in the Schaefer100 parcellation with a total of 100 cortical regions (50 regions per hemisphere)^72^ using a custom-made Matlab script using the ft_read_cifti function from the Fieldtrip toolbox^73^ to extract and average in each region. We filtered these average BOLD timeseries using a second-order Butterworth filter in the range of 0.008-0.08Hz.

### Receptor maps from PET

Receptor densities were assessed using PET tracer studies for 19 different receptors and transporters across nine neurotransmitter systems from more than 1,200 healthy participants, based on data (available at GitHub) provided by Hansen and colleagues^34^. These neurotransmitter systems include dopamine (D1, D2, DAT, noradrenaline (NET), serotonin (5HT1A, 5HT1B, 5HT2A, 5HT4, 5HT6, 5HTT), acetylcholine (A4B2, M1, VAChT), glutamate (mGLUR5, NMDA), GABA (GABA-A), histamine (H3), cannabinoid (CB1), and opioid (MOR). The volumetric PET images were aligned to the MNI-ICBM 152 nonlinear 2009 (version c, asymmetric) template, averaged across participants within each study, and then parcellated. For receptors/transporters with multiple mean images from the same tracer (5HT1B, D2, VAChT), a weighted average was applied^34^.

### Whole-brain model

The whole-brain model is using the Schaefer100 parcellation, where the local dynamics of each brain region is expressed by a Stuart-Landau oscillator (i.e., as the normal form of a supercritical Hopf bifurcation). The Hopf model has become a standard model for examining the shift from noisy to oscillatory dynamics ^74^, and they have been used to replicate key aspects of brain dynamics observed in electrophysiology^75,76^, magnetoencephalography^77^ and fMRI^78,79^.

Here, for the Schaefer100 parcellation with *N* = 100 regions, the coupling the local dynamics of *N* Stuart-Landau oscillators via the connectivity matrix ***C***, defines the whole-brain dynamics as follows

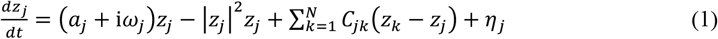

where for the oscillator in region *j*, the complex variable *z*_*j*_ denotes the state (*z*_*j*_ = *x*_*j*_ + i*y*_*j*_), *η*_*j*_ is additive uncorrelated Gaussian noise with variance *σ*^2^ (for all *j*), *ω*_*j*_ is the intrinsic node frequency, and *a*_*j*_ is the node’s bifurcation parameter. Within this model, the intrinsic frequency *ω*_*j*_ of each node is in the 0.008−0.08Hz band. The intrinsic frequencies were estimated from the data, as given by the averaged peak frequency of the narrowband blood-oxygen-level-dependent (BOLD) signals of each brain region. In **Equation 1**, *C*_*jk*_ is a specific entry in the coupling connectivity matrix ***C***, which is optimised to fit the resting-state empirical data, as detailed below. For *a*_*j*_ > 0, the local dynamics settle into a stable limit cycle, producing self-sustained oscillations with frequency *ω*_*j*_/(2*π*). For *a*_*j*_ < 0, the local dynamics present a stable spiral point, producing damped or noisy oscillations in the absence or presence of noise, respectively. The fMRI signals were modelled by the real part of the state variables, i.e., *x*_*j*_ = Real(*z*_*j*_). It has been shown that the best working point for fitting whole-brain neuroimaging dynamics is at the brink of the bifurcation, i.e. with *a*_*j*_ slightly negative but very near to zero (usually *a*_*j*_ = −0.02) ^35^. This proximity to criticality is crucial, because it allows a linearization of the dynamics ^80^, which permits an analytical solution for the functional connectivity matrix ***FC***^*model*^ (given by the Pearson correlations between all pairs of brain regions).

Briefly, we can estimate the functional correlations of the whole-brain network using a linear noise approximation (LNA). Hence, the dynamical system of *N* nodes (**Equation 1**) can be re-written in vector form as:

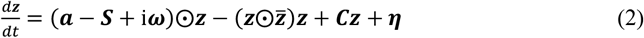

where ***z*** = [*z*_1_, …, *z*_*N*_]^*T*^, ***a*** = [*a*_1_, …, *a*_*N*_]^*T*^, ***ω*** = [*ω*_1_, …, *ω*_*N*_]^*T*^, ***η*** = [*η*_1_, …, *η*_*N*_]^*T*^ and ***S*** = [*S*_1_, …, *S*_*N*_]^*T*^ is a vector containing the strength of each node, i.e. *S*_*i*_ = ∑_*j*_ *C*_*ij*_. The superscript []^*T*^represents the transpose, 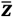 is the complex conjugate of ***z*** and ⨀ is the Hadamard element-wise product. As such, the equation describes the linear fluctuations around the fixed point ***z*** = 0, which is the solution of 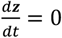. Discarding the higher-order terms 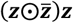 and separating the real and imaginary parts of the state variables, the evolution of the linear fluctuations follows a Langevin stochastic linear equation:

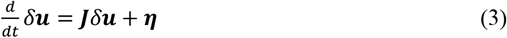

where the 2*N*-dimensional vector *δ****u*** = [*δ****x***, *δ****y***]^***T***^ = [*δx*_1_, …, *δx*_*N*_, *δy*_1_, …, *δy*_*N*_]^*T*^ contains the fluctuations of real and imaginary state variables. The 2*N* × 2*N* matrix ***J*** is the Jacobian of the system evaluated at the fixed point, which can be written as a block matrix

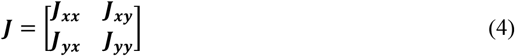

where ***J***_***xx***_, ***J***_***xy***_, ***J***_***yx***_, ***J***_***yy***_ are *N* × *N* matrices given as ***J***_***xx***_ = ***J***_***yy***_ = diag(***a*** − ***S***) + ***C*** and ***J***_***xy***_ = −***J***_***yx***_ = diag(***ω***), where diag(***v***) is the diagonal matrix whose diagonal is the vector ***v***. Please note that this linearisation is only valid if ***z*** = 0 is a stable solution of the system, that is if all eigenvalues of ***J*** have negative real parts. To compute the covariance matrix ***K*** = ⟨*δ****u****δ****u***^***T***^⟩, one can begin by writing **Equation 3** as *dδ****u*** = ***J****δ****u****dt* + *d****W***, where *d****W*** is an 2*N*-dimensional Wiener process with covariance ⟨*d****W****d****W***^***T***^⟩ = ***Q****dt*, where ***Q*** is the noise covariance matrix, which is diagonal if the noise is uncorrelated. Using Itô’s stochastic calculus, we get *d*(*δ****u****δ****u***^***T***^) = *d*(*δ****u***)*δ****u***^***T***^ + *δ****u****d*(*δ****u***^***T***^) + *d*(*δ****u***)*d*(*δ****u***^***T***^). Noting that ⟨*δ****u****d****W***^***T***^⟩ = 0, taking the expectations and keeping terms to first order in the differential *dt*, we obtain:

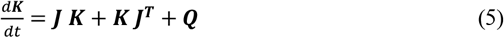

Hence, the stationary covariances (for which 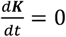) can be obtained by solving the following analytic equation:

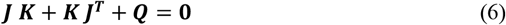

**Equation 6** is a Lyapunov equation that can be solved using the eigen-decomposition of the Jacobian matrix^81^. We obtained the simulated functional connectivity matrix ***FC***^*model*^ from the first *N* rows and columns of the covariance ***K***, which corresponds to the real part of the dynamics, and thus the BOLD fMRI signal.

### Optimizing the resting state generative effective connectivity (RS-GEC)

For defining the RS-GEC of a whole-brain model, we first fit the model to the empirical resting state data for each participant. For this, we used a noise *η*_*j*_ with variance *σ*^2^ = 0.02 homogeneous for all regions in the Schaefer100 parcellation. In order to create the resting state generative effective connectivity (RS-GEC), we used a pseudo-gradient procedure to optimise the initial coupling connectivity matrix ***C***, derived from the anatomical structural connectivity which was then iteratively used to create the RS-GEC.

Specifically, we iteratively compared the output of the model with the empirical measures of the functional correlation matrix (***FC***^*empirical*^), i.e., the normalised covariance matrix of the functional neuroimaging data. Furthermore, we also compared the output of the model with the normalized *τ* time-shifted covariances (***FS***^*empirical*^(*τ*)). These normalised time-shifted covariance matrices are generated by taking the shifted covariance matrix ***KS***^*empirical*^(*τ*) and dividing each pair (*i, j*) by 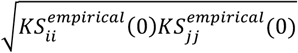. Note that these normalised time-shifted covariances break the symmetry of the couplings and thus improve the level of fitting ^82^. Using a heuristic pseudo-gradient algorithm, we proceeded to update the ***C*** until the fit is fully optimised. More specifically, the updating uses the following form:

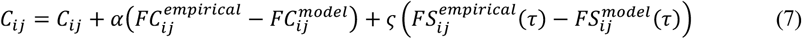

where 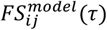 is defined similar to 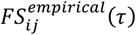. In other words, the updating is given by the first *N* rows and columns of the simulated *τ* time-shifted covariances ***KS***^*model*^(*τ*) normalised by dividing each pair (*i, j*) by 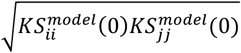, where ***KS***^*model*^(*τ*) is the shifted generative covariance matrix computed as follows:

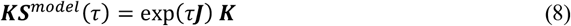

Please note that ***KS***^*model*^(0) = ***K***. The model was then run repeatedly with the updated ***C*** until the fit converges towards a stable value. As stated above, we initialised ***C*** using the average anatomical connectivity (obtained with probabilistic tractography from dMRI, see *Methods* above) and only update known existing connections from this matrix (in either hemisphere). There is one exception, however, to this rule which is that the algorithm also updates homologue connections between the same regions in either hemisphere, given that tractography is known to be less accurate when accounting for this connectivity.

For the Stuart-Landau model, we used *α* = *ς* = 0.0004 and continue until the algorithm converges. For each iteration we compute the model results as the average over as many simulations as there are participants. We applied this procedure to fit a whole-brain model to resting state data for each individual which generated 971 individual RS-GEC, which were subsequently used in the framework described as follows.

### Local dynamics optimised with neurotransmission modulation (NEMO)

In order to make it possible for a whole-brain model with the fixed RS-GEC architecture to compute the different task dynamics, we used neurotransmission modulation (NEMO) of the local regional dynamics. They were optimised to modulate the RS-GEC model to describe the task functional connectivity (TASK-FC) for each individual. This was possibly by optimising the local regional dynamics through finding the optimal noise values ***η*** = [*η*_1_, …, *η*_*N*_]^*T*^ for each region such that the model-generated and empirical TASK-FC are maximally correlated using the Structural Similarity Index (SSIM). The noise *η*_*i*_ in each region *i* (with *i* from 1 to *N*) is given by

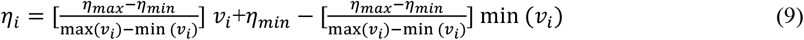

where *v*_*i*_ is the weighted sum of the 19 neurotransmitter maps given by

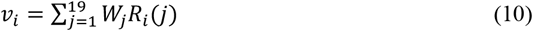

where *W*_*j*_ are the weighting parameters to be optimised constrained to the range [−1 1]. Please note that the noise *η*_*i*_ values were constrained between a minimum and maximum values, *η*_*min*_ and *η*_*max*_ which were also optimized constrained to the range [0.001 0.04]. Accordingly, the final noise *η*_*i*_ utilized in the model is bounded between *η*_*min*_ and *η*_*max*_.

Importantly, the neurotransmitter maps were normalised following the procedure of Fulcher and Fornito^83^. The maps are z-scored across the 100 regions in the Schaefer100 parcellation and then non-linearly normalised by a sigmoid function to reduce the influence of outliers:

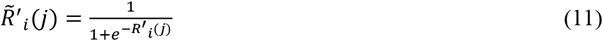

where *R*^*′*^_*i*_(*j*) signifies the receptor *j* in the region *i* from the original neurotransmitter maps provided by Hansen and colleagues ^34^.

Hence, the normalised neurotransmitter maps are given by

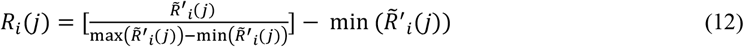

In order to produce the optimised weights, we employed MATLAB’s global optimization tool known as the particle swarm optimiser. This method is akin to genetic algorithms which are population-based approaches that have been extensively applied to complex optimisation tasks. The algorithm simulates a swarm of particles—comparable to starlings exploring a space—which traverse the parameter search domain^84–86^. The swarm algorithm adaptively guides the particles’ movement to effectively locate the global optimum. Here, we used the swarm optimiser with 21 parameters (in addition to the 19 *W*_*j*_ weights, we also include the extrema of the noise *η*_*min*_ and *η*_*max*_) in order to maximise the SSIM between the empirical and model-generated FC matrices.

### Brain Computability

For each participant, brain computability is defined as the average performance of the individualised NEMO whole-brain models across the seven tasks. For each task, we first quantified the level of performance, called ‘brain computation’ using SSIM, which measures the similarity between the empirical task-based functional connectivity (FC) matrix and the FC matrix generated by the NEMO whole-brain model for this task. For each task, we established a normalised reference point, by dividing these SSIM values by the SSIM between the empirical resting-state FC and the RS-GEC model-generated output (i.e., the original resting-state model without neurotransmitter modulation). This normalisation was performed at the individual or group level, depending on the specific experimental setup, as detailed in the *Results*.

### Measuring the g-factor

The *g*-factor was computed using a matlab adaptation of the procedure described by Dubois and colleagues^48^ to perform factor analysis of the scores on ten relevant cognitive scores from the behavioural psychometric battery used to assess each HCP individuals. This procedure derives the *g*-factor measure of intelligence, which is a standard used in the field of intelligence research.

### Defining global brain connectivity (GBC)

The global brain connectivity (GBC) can be computed from the functional connectivity matrix **FC** as follows

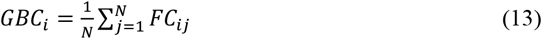

### Working point analysis

For the working point analysis in **Figure 5A**, we used the average RS-GEC across over all individuals and multiplied this by the global coupling factor, *G*. This defines the working point of the whole-brain model as *G*=1. We calculated brain computability of the NEMO whole-brain model for each point of *G*=[0.2 1.8] (in steps of 0.1) which was run 10 times with averaged normalised weights across all individuals for each of the seven tasks.

### Measure of hierarchical organisation

For measuring the hierarchical organisation of the brain networks, we adapted the hierarchy measures of directedness and trophic levels in directed networks ^46,87^. We generate both the hierarchical node level information, *trophic level*, and the global information, *directedness* (or trophic coherence), which are applied the directed graphs obtained from the GEC matrices, the individualised optimised RS-GEC matrix.

As stated above, the RS-GEC (***C***) matrix defines a graph of *N* nodes connected by weighted edges determined by its elements. For each node *n* in the graph, we introduce the concepts of in-weight 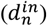 and out-weight 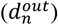 defined as follows:

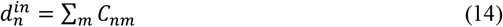

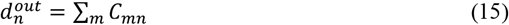

We define the total weight of node *n* as *u*_*n*_ by

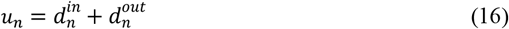

In addition, we define the imbalance for node *n* as *v*_*n*_, representing the difference between the flow into and out of the node by

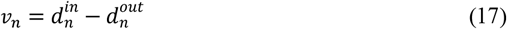

The (weighted) graph-Laplacian operator **Λ** on vectors ***h***, is given by

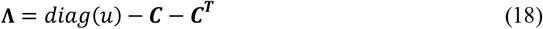

Consequently, the enhanced concept of trophic level corresponds to the solution ***h*** of the linear system of equations

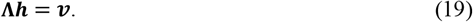

Here each component of the vector ***h*** corresponds to the trophic level in a brain region. Importantly, while the operator **Λ** is symmetric, the asymmetry of the network is evident in the imbalance vector ***v***.

Once the hierarchy level *h* has been established, we can assess the network’s global directionality by computing its directedness (or trophic coherence) using following equation:

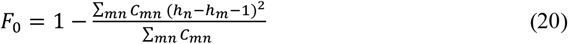

A network is considered maximally coherent when *F*_0_=1, whereas it is regarded as incoherent when *F*_0_=0. The trophic coherence is a graph theoretical measure of hierarchical organisation. Elevated values of trophic coherence indicate a greater degree of hierarchical organisation.

### Measure of non-equilibrium

Measuring the level of non-equilibrium was performed by using a framework for measuring the violation of the fluctuation-dissipation theorem (FDT)^50^. All details are included in the original paper where it was shown that the deviation from the FDT for a given brain region *i* when a perturbation is applied to brain region *j* is given by:

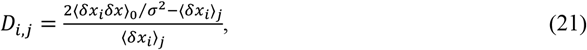

where the term 2/*σ*^2^ plays the role of the inverse temperature 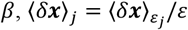, i.e. the real part of ⟨*δ****u***⟩_*j*_ and the covariance ⟨*δx*_*i*_*δx*⟩_0_ is derived from ***KS***^*model*^. For numerical reasons, we quantify the system-wide effect of perturbing the component *j* by averaging the numerator and denominator over the regions; i.e.,

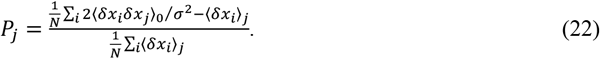

Here, vector *P* defines a *perturbability map* over all brain regions in a given brain state. For each participant, the level of non-equilibrium 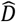 is computed by averaging the deviation from the FDT over all possible perturbations; i.e.,

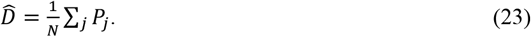

## Data availability

The neuroimaging data are freely available from HCP.

## Code availability

The code used to run the analysis from the HCP data is available on GitHub (https://github.com/decolab/braincomp).

## Acknowledgements

G.D. is supported by Grant PID2022-136216NB-I00 funded by MICIU/AEI/10.13039/501100011033 and by “ERDF A way of making Europe”, ERDF, EU, Project NEurological MEchanismS of Injury, and Sleep-like cellular dynamics (NEMESIS) (ref. 101071900) funded by the EU ERC Synergy Horizon Europe, and AGAUR research support grant (ref. 2021 SGR 00917) funded by the Department of Research and Universities of the Generalitat of Catalunya. Y.S.P. is supported by was supported by the project NEurological MEchanismS of Injury, and Sleep-like cellular dynamics (NEMESIS) (ref. 101071900) funded by the EU ERC Synergy Horizon Europe. M.L.K. is supported by the Centre for Eudaimonia and Human Flourishing (funded by the Pettit and Carlsberg Foundations) and Center for Music in the Brain (funded by the Danish National Research Foundation, DNRF117). The funders had no role in study design, data collection and analysis, decision to publish or preparation of the manuscript.

## Author Contributions

All the authors (GD, YSP, JV, AL & MLK) designed the study, developed the methods, performed the analyses, and wrote and edited the manuscript. All the authors edited the manuscript.

